# Task learning is subserved by a domain-general brain network

**DOI:** 10.1101/2022.12.07.519504

**Authors:** Jiwon Yeon, Alina Sue Larson, Dobromir Rahnev, Mark D’Esposito

## Abstract

One of the most important human faculties is the ability to acquire not just new memories but the capacity to perform entirely new tasks. However, little is known about the brain mechanisms underlying the learning of novel tasks. Specifically, it is unclear to what extent learning of different tasks depends on domain-general and/or domain-specific brain mechanisms. Here human subjects (N=45) learned to perform six new tasks while undergoing functional MRI. The different tasks required the engagement of perceptual, motor, and various cognitive processes (attention, expectation, speed-accuracy tradeoff, and metacognition). We found that a bilateral frontoparietal network was more active during the initial compared to the later stages of task learning, and that this effect was stronger for task variants requiring more new learning. Critically, the same frontoparietal network was engaged by all six tasks, demonstrating its domain generality. Finally, although task learning decreased the overall activity in the frontoparietal network, it increased the connectivity strength between the different nodes of that network. These results demonstrate the existence of a domain-general brain network whose activity and connectivity reflect learning for a wide variety of new tasks, and thus may underlie the human capacity for acquiring new abilities.

## Introduction

Humans have the remarkable ability to learn completely new skills. Learning is most vigorous in childhood when people master various skills such as reading, writing, arithmetic, sports, and self-control (Piaget, 1970). Yet, neurologically healthy humans never lose their ability to acquire new skills, and adults routinely learn new languages, sports, job skills, and social habits. However, it remains unclear what brain mechanisms underlie our ability to learn such a diverse range of new skills. Specifically, a central problem is the extent to which learning across a variety of domains relies on common neural mechanisms vs. domain-specific mechanisms that are different for each task.

Previous research has focused primarily on the neural correlates of learning specific stimuli. For example, studies investigating perceptual learning typically compare brain activity associated with highly trained versus untrained sensory stimuli (Antzoulatos & Miller, 2011; Rainer et al., 2004; Yang & Maunsell, 2004), studies on motor learning compare familiar and novel movements (Bassett et al., 2015; Grafton et al., 2002; Houweling et al., 2008; Musall et al., 2019), studies on classical and operant conditioning compare rewarded and unrewarded stimuli (Baeuchl et al., 2020; Serences, 2008; Shuler & Bear, 2006; Summerfield & Koechlin, 2010), and studies on repetition suppression compare repeated and non-repeated stimulus presentations (Henson et al., 2000; Wig et al., 2005). In all of these cases, what is being investigated is how a learned stimulus or action differs from a novel stimulus or action. The same limitation also applies to other common designs, such as to studies that compare stimulus-evoked brain activity before and after learning (Deuker et al., 2013; Peter et al., 2021; Serences et al., 2009; Stauch et al., 2021; Utzerath et al., 2017). However, despite the tremendous progress made with these approaches, this line of research tells us very little about the brain mechanisms that underlie our ability to learn a new *task* rather than a new *stimulus.*

Not surprisingly, previous research focusing on comparing learned versus novel stimuli has mostly found that domain-specific mechanisms underlie each area of learning. After extensive training, visual stimuli are processed differently in the visual cortex (Deuker et al., 2013; Rainer et al., 2004; Serences, 2008; Serences et al., 2009; Shuler & Bear, 2006; Utzerath et al., 2017), motor actions are generated differently in the motor cortex (Bassett et al., 2015; Grafton et al., 2002; Houweling et al., 2008; Musall et al., 2019), and rewarded stimuli are processed differently in reward circuits of the brain (Baeuchl et al., 2020; Serences, 2008; Shuler & Bear, 2006; Summerfield & Koechlin, 2010). Based on this research, it is perhaps natural to hypothesize that the ability to learn completely novel tasks that rely on different perceptual, motor, and cognitive processes would rely on domain-specific brain areas. However, an alternative possibility is that learning novel tasks depends on domain-general mechanisms and that domain-specific brain areas become important primarily when learning specific stimuli, but not the structure associated with a new task.

Several studies have investigated task learning for tasks that share a common meta-structure. For example, one approach introduced by Cole et al. (2010) involves constructing 64 different tasks specified using combinations of judging one of four sensory properties (green/loud/soft/sweet), applying one of four logic operations (same/just one/second/not second), and executing one of four motor responses (left/right index/middle finger). Thus, the instructions for one of the 64 tasks would be “If the answer to ‘Is it GREEN?’ is the SAME for both words, press your LEFT INDEX finger.” The task consists of observing two words (e.g., “gecko” and “leaf”) and making an appropriate response. Several other designs for novel task learning use simpler task structures that consist of learning different stimulus-response pairings and implementing them either immediately (Hartstra et al., 2011; Ruge & Wolfensteller, 2010) or after a delay (Meiran et al., 2015a, 2015b). These studies generally found that novel task variants engaged frontal and parietal regions more strongly than previously practiced variants. However, in all of these studies, the different tasks are always specific instances of the same overall task structure, thus making it difficult to establish whether learning tasks from different domains rely on domain-general or domain-specific mechanisms.

Here we adjudicated between these two possibilities. Human subjects (N=45) learned to perform six new tasks that rely on perception, motor, or various cognitive processes (attention, expectation, speed-accuracy tradeoff, and metacognition). Critically, subjects were first introduced to each new task inside an MRI scanner, which allowed us to examine their brain activity during the process of learning each new task. Each task had three variants that differed in several dimensions, while remaining within the same task domain. Importantly, each variant consisted of two blocks, and we compared the brain activity during the first block when substantial new task learning is necessary to the second block when relatively less new task learning is taking place. We observed that a domain-general frontoparietal network subserved learning across all six tasks. This network showed higher activity during the initial stages of learning and stronger connectivity in the later stages of learning. These results demonstrate the existence of a domain-general network that underlies task learning.

## Results

We investigated whether the neural substrates underlying the learning of new tasks is domain-general or domain-specific. Subjects performed one perceptual, four cognitive, and one motor task while we collected fMRI data (**Figure 1**). The perception task was always performed first, the motor task was always performed last, and the remaining tasks were performed in randomized order. To avoid the introduction of new perceptual learning with each task, the cognitive tasks used the same grating stimuli from the perception task. Each task consisted of three variants to ensure the need for constant learning of new task features. Each variant included two blocks of 30 trials with the first block naturally requiring more novel learning than the second block.

**Figure 1.**
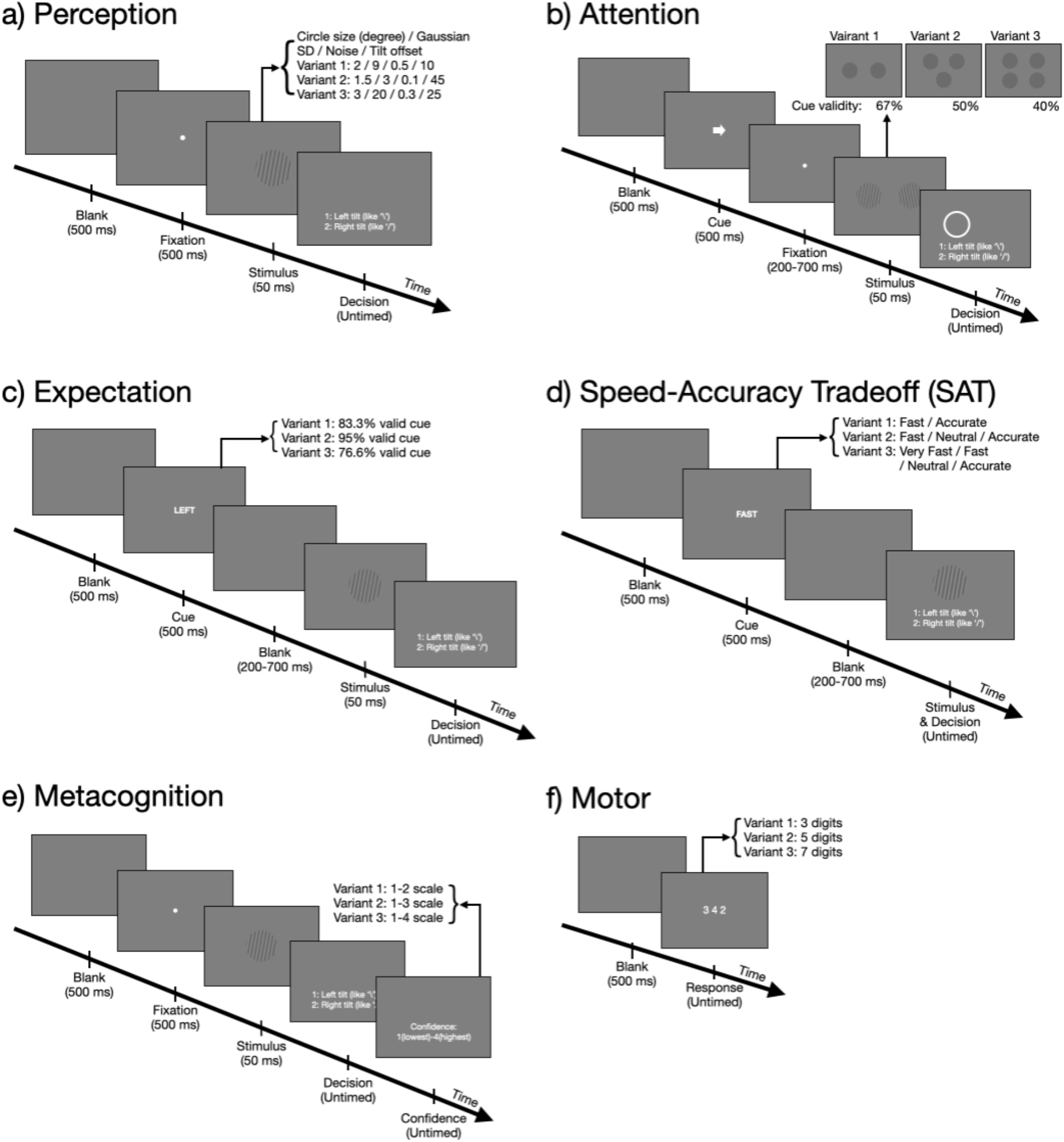
Tasks. Subjects completed a perception task, followed by four cognitive tasks (requiring attention, expectation, speed-accuracy trade-off, and metacognition) in a randomized order, and a motor task that always came last. Each task had three variants shown on the top-right of each panel. (a) In the perception task, subjects judged the orientation (left vs. right) of a grating. The three variants manipulated the size, noise level, contrast level, and orientation of the grating. The same parameters of the last variant were used for generating the stimuli in all cognitive tasks (the only exception is that the attention task used higher contrast). (b) In the attention task, a pre cue indicated the location that subjects should attend to. Subjects reported the stimulus orientation at a post-cued location. The three variants differed in the number of gratings presented and corresponding pre-cue validity. (c) In the expectation task, a cue indicated the likely orientation of the upcoming stimulus. The three variants differed in the predictiveness of the cue. (d) In the speed-accuracy trade-off task, a cue indicated the speed with which subjects should make the orientation discrimination response. The three variants differed in the number of speed stress levels. (e) In the metacognition task, subjects provided a confidence rating regarding the accuracy of their decision. The three variants differed in the confidence scale used. (f) In the motor task, subjects typed as quickly as possible the digits (chosen among the digits 1 to 4) presented on the screen using a button box. The three variants differed in the number of digits shown.

### Behavioral effects

We first confirmed that subjects performed all behavioral tasks as instructed. We observed adequate performance across all tasks (accuracy = 70.6%, 59.3%, 77%, 77.7%, 69.7%, and 85.9% for the perception, attention, expectation, SAT, metacognition, and motor tasks, respectively). In addition, we confirmed that subjects were more accurate for valid cues in the attention task (**Supplementary Figure 1**), preferentially chose the expected stimulus in the expectation task (**Supplementary Figure 2**), followed the speed/accuracy instructions in the SAT task (**Supplementary Figure 3**), and exhibited higher accuracy for trials with higher confidence in the metacognition task (**Supplementary Figure 4**).

We then examined whether there was behavioral evidence for a learning effect when comparing Blocks 1 and 2 of each task variant. We found that subjects exhibited substantially reduced reaction times (RTs) for Block 2 compared to Block 1. This effect was most pronounced for the first task variant (RT difference = 220 ms; *t*(44) = 9.82, *p* = 1.16 x 10^-12^; **Figure 2**) but was also present for the second (RT difference = 124 ms; *t*(44) = 5.20, *p* = 4.91 x 10^-6^) and third (RT difference = 74 ms; *t*(44) = 4.09, *p* = 1.78 x 10^-4^) task variants. Moreover, RT difference between Blocks 1 and 2 was larger for the first variant compared to both the second (*t*(44) = 3.92, *p* = 3.02 x 10^-4^) and the third (*t*(44) = 6.92, *p* = 1.51 x 10^-8^) variants. The RT difference between the second and the third variants was only marginally significant (*t*(44) = 1.84, *p* = .074). These results provide strong evidence for behavioral learning in all three task variants, with the learning being greatest in the first variant.

**Figure 2.**
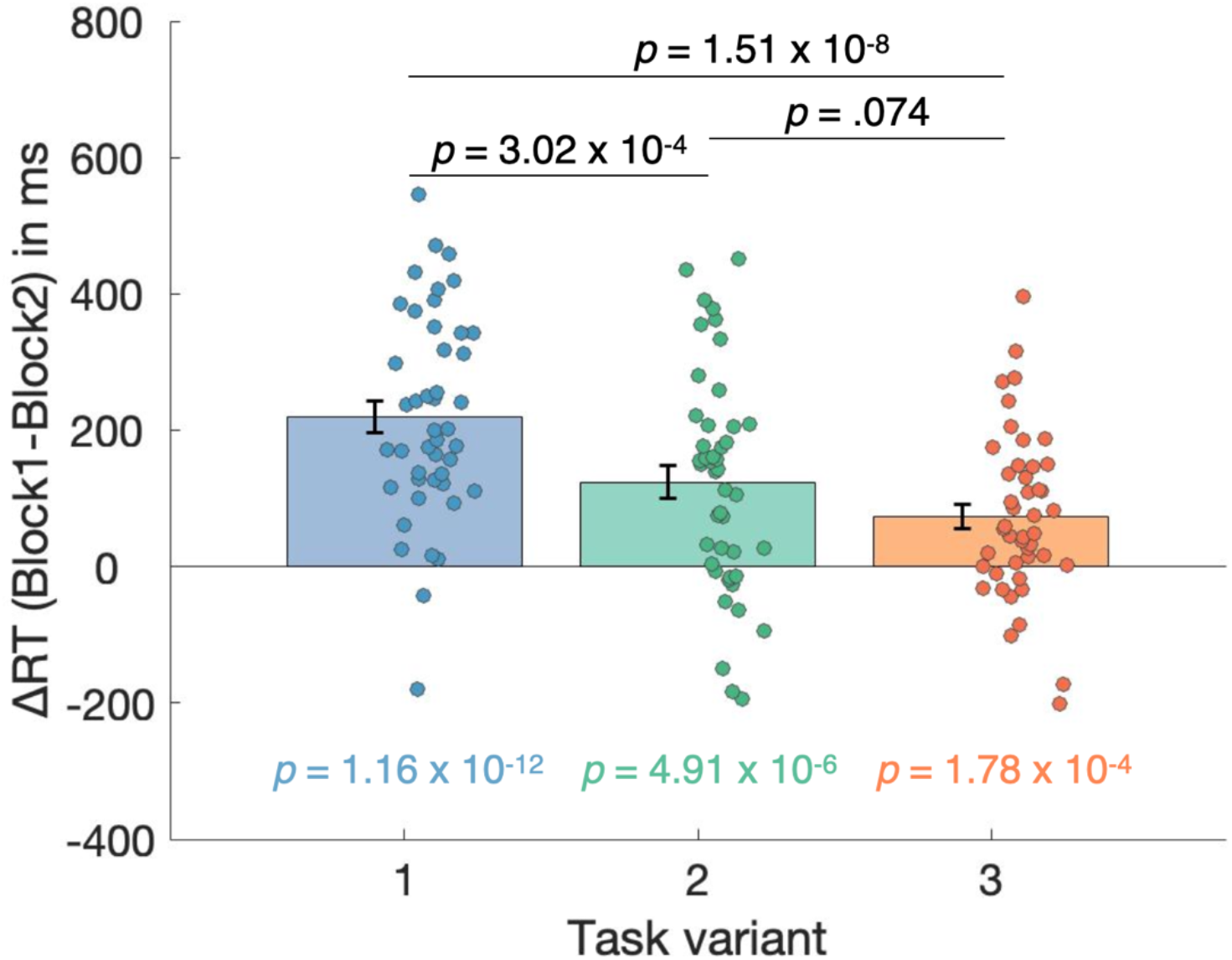
RT difference between Blocks 1 and 2 for each task variant. We computed the RT difference between Blocks 1 and 2 for each of the three task variants across all six tasks. The *p*-values below the plots show the results of one-sample *t*-tests comparing the RT difference for each of the three task variants against zero. The *p*-values above the plots show the results of paired *t*-tests comparing the RT differences between different task variants. The results suggest that learning was greatest for the first task variant but remained significant for the second and third variants. Each dot represents the average RT across all six tasks for a single subject. Error bars indicate SEM.

### Task learning is reflected in frontal, parietal, and cerebellar activation increases

The behavioral analyses confirmed that substantial task learning took place during Block 1 (as reflected by the faster RTs on Block 2 compared to Block 1). Therefore, we investigated the neural correlates of task learning by determining the brain areas that show larger activations for Block 1 than Block 2. We applied family-wise error (FWE) correction at *p* < .05 and searched for clusters of at least 150 voxels. The results showed bilateral activation in the inferior frontal gyrus (IFG), bilateral activation in the intraparietal sulcus (IPS), and left cerebellum activation (**Figure 3**, **Table 1**). No voxels survived the same threshold for the opposite comparison (Block 2 > Block 1).

**Figure 3.**
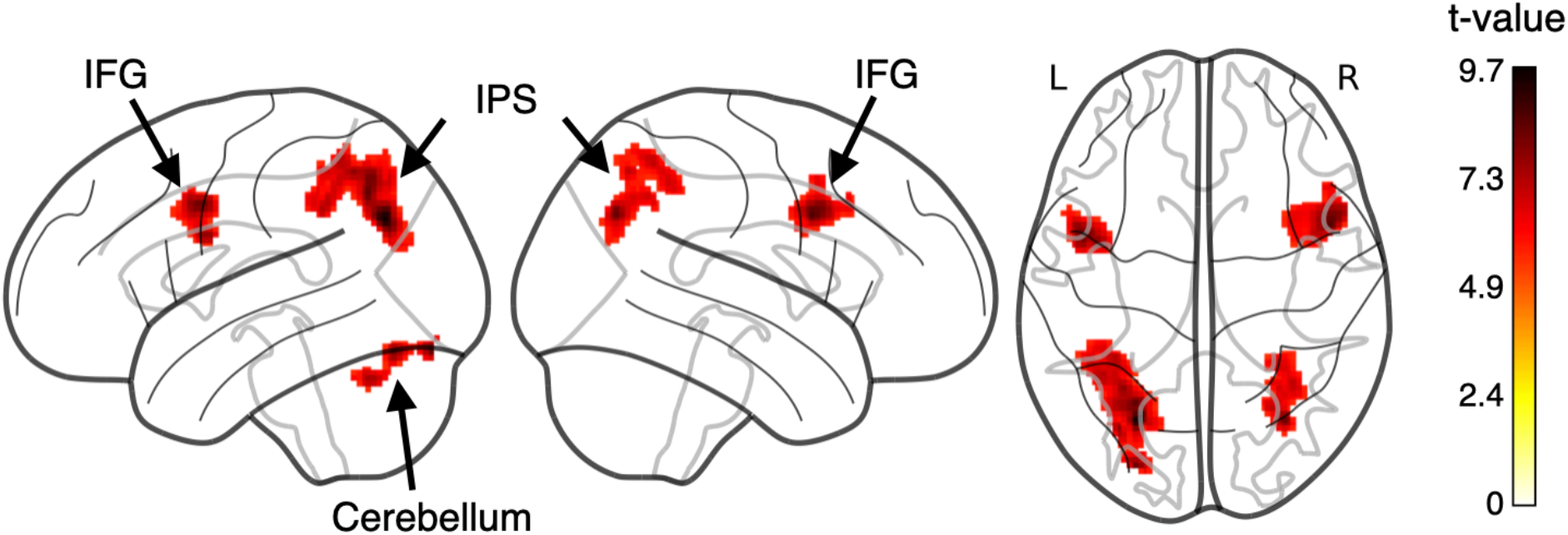
Task learning is reflected in frontal, parietal, and cerebellar regions. Brain regions with greater activation for Block 1 compared to Block 2 across all six tasks. Five clusters emerged in bilateral inferior frontal gyrus (IFG), bilateral intraparietal sulcus (IPS), and left cerebellum (*p* < .05 FWE corrected, cluster size ≥ 150). Colors indicate *t*-values.

**Table 1.**
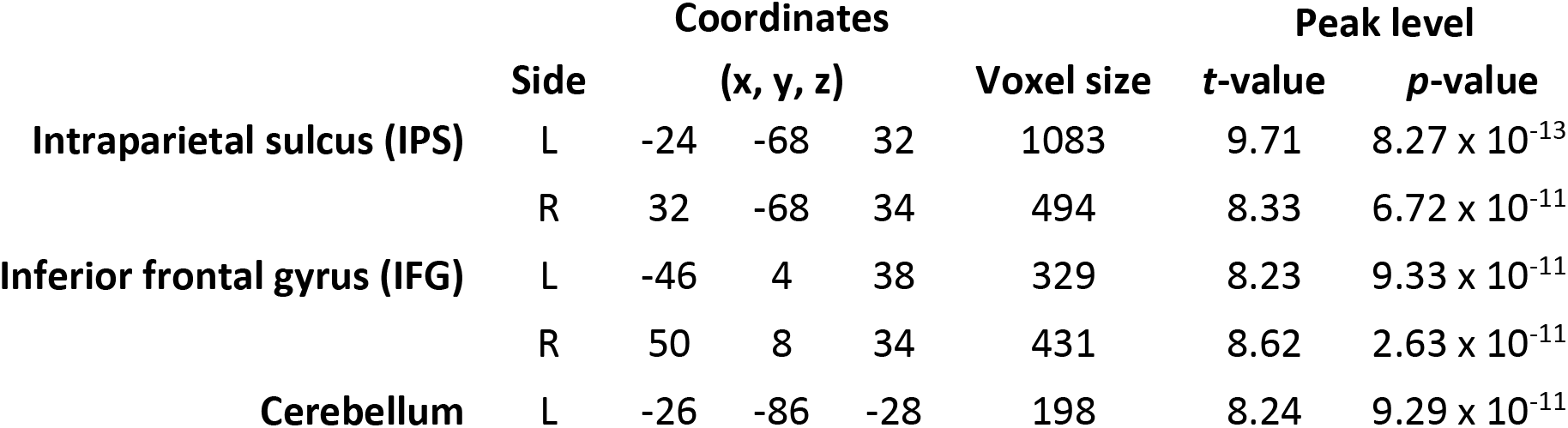
Localization of activations for the contrast Block 1 > Block 2. Coordinates correspond to the standard MNI brain.

The behavioral results demonstrated that learning was greatest for the first variant of each task and then monotonically decreased for the second and third variants (**Figure 2**). Therefore, we examined whether the difference in activation between Blocks 1 and 2 in these five clusters also peaked for the first variant and decreased for later variants. Qualitatively, the comparison of Blocks 1 and 2 produced large swaths of activity in the first variant, less activity in the second variant, and even less activity in the third variant (**Figure 4a**). We quantified this effect by examining the activations for each task variant specifically for the five regions from Figure 3, which we defined as regions-of-interest (ROIs). We found that all four frontoparietal ROIs showed a monotonic decrease in activation difference between Blocks 1 and 2 *(p* < .001 for all pairwise comparisons between variant 1 and variant 3; **Figure 4b**). While the cerebellar ROI also showed a pattern of monotonic decreases, none of the pairwise comparisons between the three variants were significant (all *p*’s > .1). Thus, even though the bilateral IFG and IPS ROIs were defined using a contrast that was independent of variant order, each of them showed a neural effect that mimicked the corresponding behavioral learning effect. An exploratory analysis also revealed a significantly positive correlation for the first variant only between (i) the RT difference between Blocks 1 and 2 averaged across six tasks and (ii) the brain activation difference between Blocks 1 and 2 averaged across six tasks and all five ROIs (**Supplementary Figure 5**), though such across-subject brain-behavior correlations should be interpreted with caution (Marek et al., 2022). Overall, these results strongly suggest that learning of these tasks is reflected in the activity in bilateral IFG, bilateral IPS, and, to a lesser extent, in left cerebellum.

**Figure 4.**
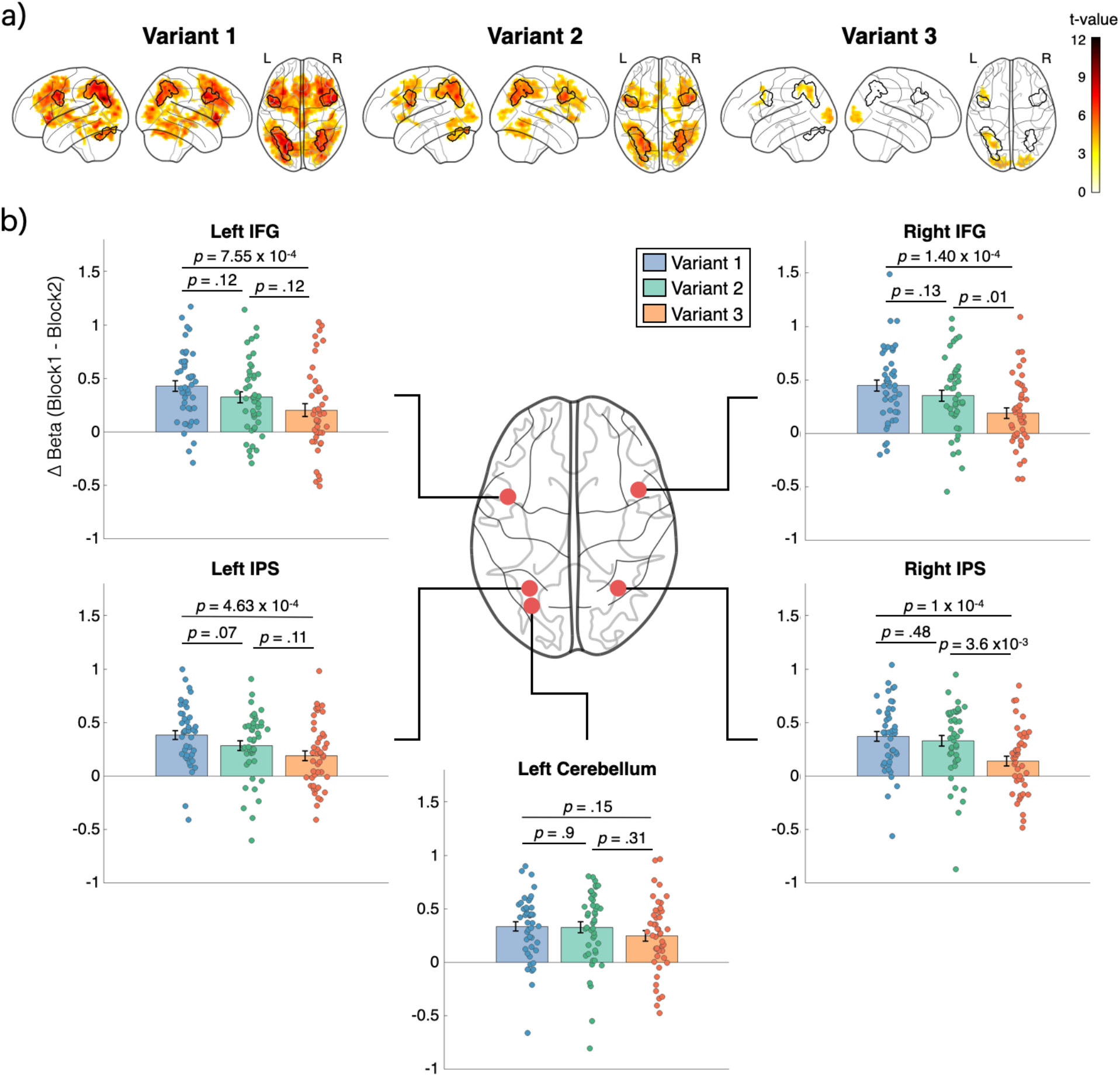
Brain activations for each of the three task variants. (a) Brain regions with greater activation for Block 1 compared to Block 2 across all six tasks displayed separately for each of the three task variants. For display purposes, the figures are thresholded at *p* < .001 uncorrected, cluster size ≥ 150. Brain activations are clearly strongest for the first task variant and weakest for the third. Black borders indicate the five regions from Figure 2, and colors indicate *t*-values. (b) Beta values difference between Blocks 1 and 2 for each of the five ROIs from Figure 2. Each ROI shows a decreasing trend such that the activation difference between Blocks 1 and 2 is greatest in the first task variant and smallest in the third, but the comparison is only significant in the four frontoparietal regions. Error bars indicate SEM. IFG, inferior frontal gyrus; IPS, intraparietal sulcus.

### Task learning effects in frontal, parietal, and cerebellar regions are domain-general

The results above clearly demonstrate that the frontal, parietal, and cerebellar brain regions reflect task learning, but do not clarify whether these regions are domain-general. Specifically, it is possible that each task exhibits strong domain-specific effects that are obscured by examining all tasks together. To explore this possibility, we examined the contrast between Blocks 1 and 2 separately for each of the six tasks. Previous imaging studies have demonstrated that visual perception tasks activate the visual cortex (Grill-Spector & Malach, 2004; Heeger, 1999), that top-down attention tasks activate the dorsal attention network (Corbetta & Shulman, 2002; Rahnev et al., 2012), that expectation tasks activate the dorsolateral prefrontal cortex, intraparietal sulcus, and medial temporal cortex (Rahnev, Lau, et al., 2011), that speed-accuracy trade-off tasks activate the supplementary motor area (Forstmann et al., 2008; Spieser et al., 2017), that metacognitive tasks activate the anterior prefrontal cortex, dorsolateral prefrontal cortex, and anterior cingulate (Rahnev et al., 2016; Shekhar & Rahnev, 2018; Yeon et al., 2020), and that motor control tasks activate motor and premotor cortices (Laut Ebbesen & Brecht, 2017; Svoboda & Li, 2018). A domain-specific account of task learning would predict that the same areas involved in the execution of each task would also be activated when learning the corresponding task. However, we found that the learning-related activations for the individual tasks were remarkably similar such that all six tasks showed activations that overlapped substantially with the across-task results (**Figure 5**), with the only exception of a prominent visual cortex activation for the perception task. Therefore, these results do not support the domain-specific account of task learning.

**Figure 5.**
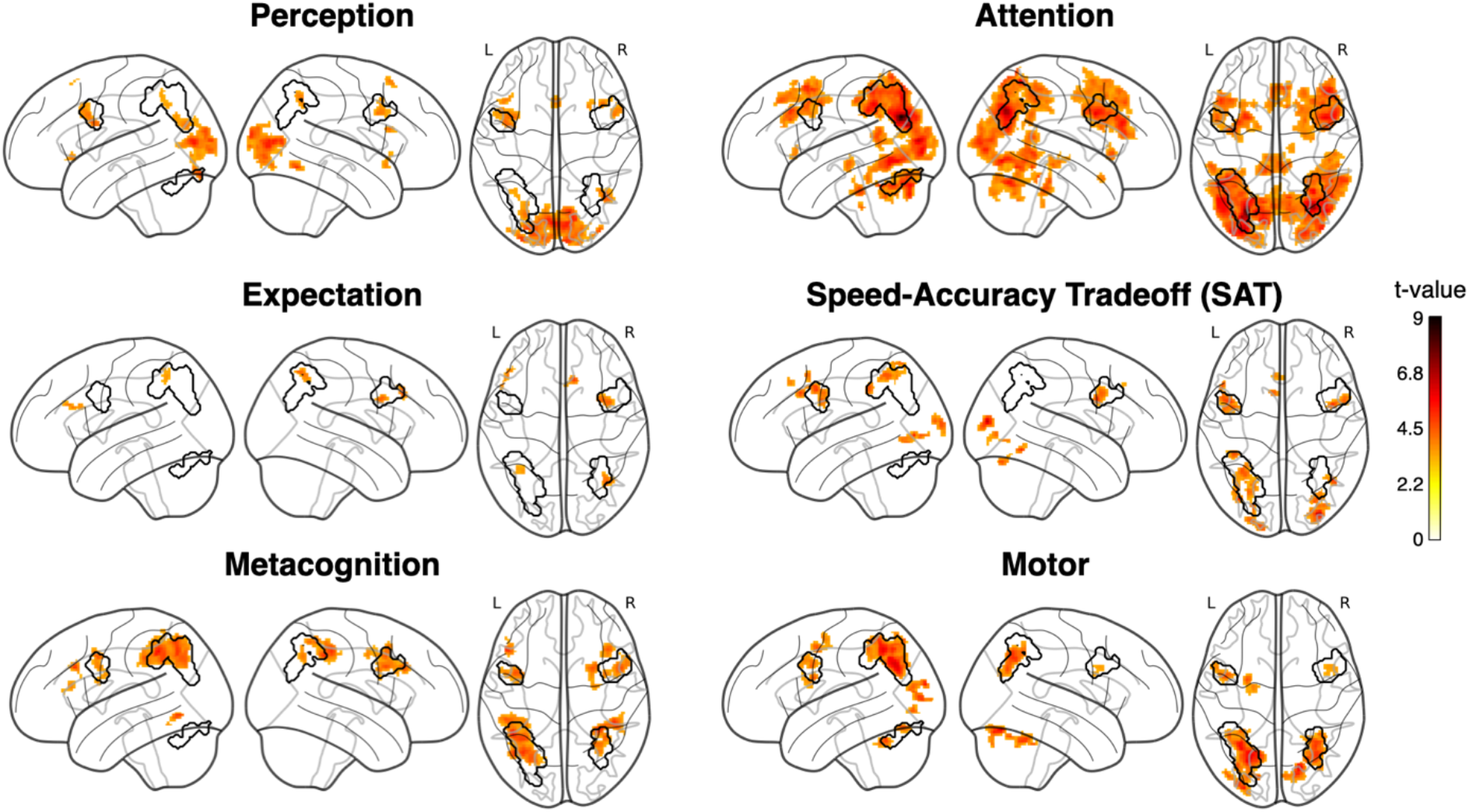
Similar brain correlates of task learning for each of the six tasks. Brain regions with greater activity for Block 1 compared to Block 2 for each task. For display purposes, the figures are thresholded at *p* < .001 uncorrected with cluster size ≥ 30. Despite notable variability in the strength of the effects across tasks, a qualitatively similar activity pattern appears across all six tasks. No domain-specific activations were found for any of the tasks, with the exception of a prominent visual cortex activation cluster in the perception task. Black borders indicate the five regions from Figure 2, colors indicate *t*-values.

These results qualitatively support the existence of a domain-general mechanism for task learning, but do not establish that all five ROIs are involved in each of the six tasks. To explore this issue, we performed a more direct test of whether each of the five areas above (bilateral IFG, bilateral IPS, and left cerebellum) were activated for all or just for a subset of the tasks investigated here. For each task, we defined the five regions as ROIs based exclusively on the data from the remaining five tasks. For example, the ROIs used for the perceptual task were defined solely based on the attention, expectation, SAT, metacognition, and motor tasks. This procedure avoided “double-dipping” where the same data are used both to define and test an ROI (Kriegeskorte et al., 2009). We then tested whether activity in these ROIs was stronger for Block 1 than Block 2 in the left-out task. We found that this was the case for 29 of the 30 tests (all *p’s* < .05), with the only exception being the left cerebellum ROI in the expectation task (**Figure 6**). These results confirm that the activity found within the frontal, parietal, and cerebellar ROIs associated with task learning are indeed domain-general and not driven by only a subset of tasks.

**Figure 6.**
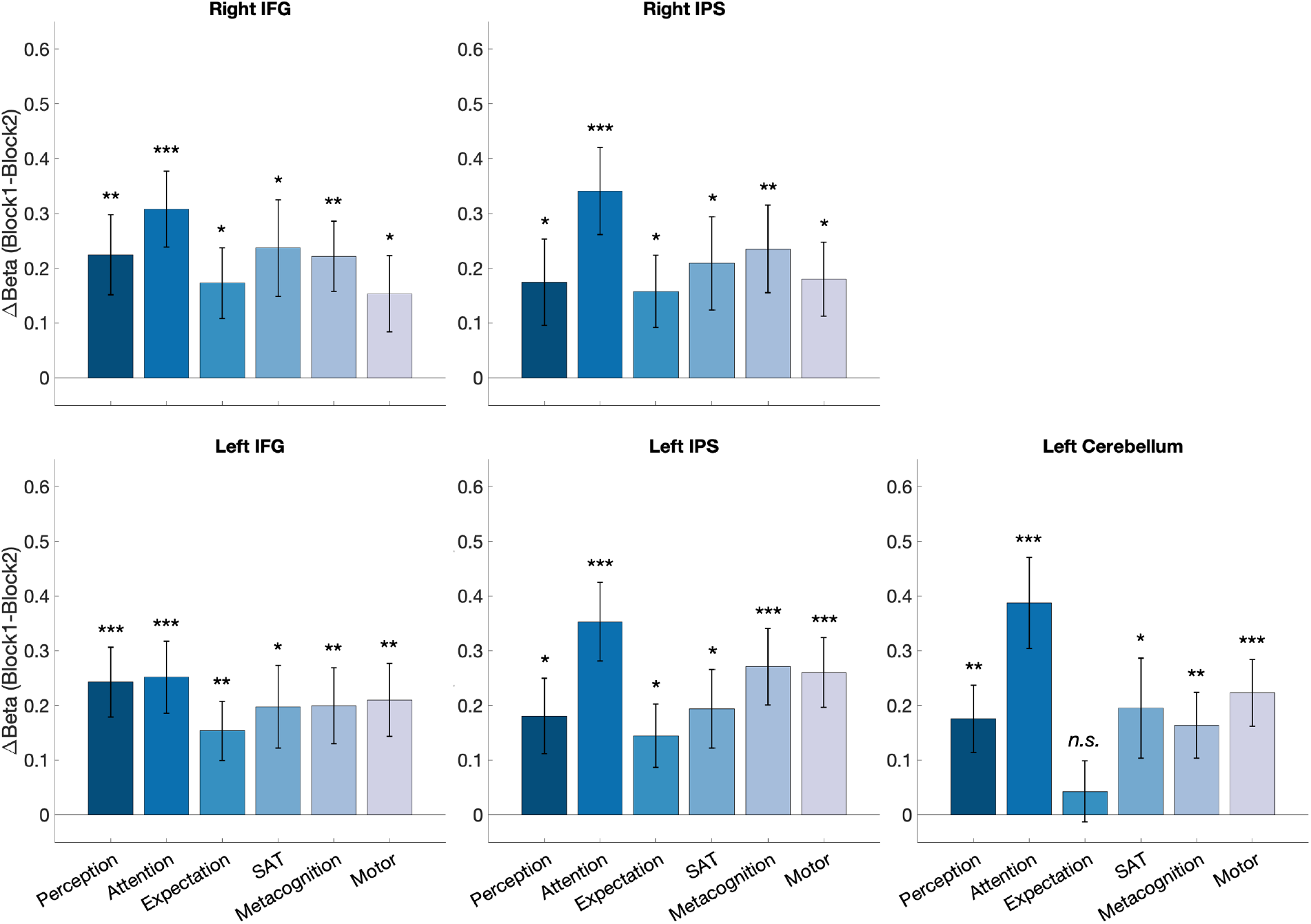
Domain generality of the activations in the frontal, parietal, and cerebellar regions. To assess the domain generality of the five ROIs (left and right IFG, left and right IPS, and left cerebellum), we defined each ROI based on the activations in five tasks and examined the activations in the left-out task. Except for the left cerebellum ROI in the expectation task, all ROIs showed a significantly larger activation for Block 1 compared to Block 2 (all *p’s* < .05). These results establish the domain generality of each of the five ROIs by demonstrating that each ROI reflects the learning on all six tasks (or five tasks in the case of left cerebellum). Error bars indicate SEM.

### Learning increases functional connectivity between frontoparietal regions

The results above establish that task learning is subserved by domain-general activations in bilateral IFG, bilateral IPS, and left cerebellum. Yet, it is unclear whether these five regions form a single network, and whether learning changes the way they communicate with each other. Here we examined these questions by investigating the functional connectivity between all five regions separately for Blocks 1 and 2. We found strong functional connectivity within the bilateral frontoparietal regions (all *r*-values between .453 and .777; **Figure 7a,b**) but much weaker functional connectivity between the left cerebellar region and the frontoparietal areas (all *r*-values between .107 and .234). In fact, each of the six within-frontoparietal network *r*values was significantly higher than each of the four cerebellum-to-frontoparietal *r*-values for both Blocks 1 and 2 (*p* < .00001 for all 48 tests). Thus, it appears that the four bilateral frontoparietal regions form a single network that the left cerebellar region is not a part of.

Critically, we examined the changes of functional connectivity between Blocks 1 and 2. We found that functional connectivity between the four frontoparietal regions was significantly higher in Block 2 (average *r* = .618) than Block 1 (average *r* = .582; *t*(44) = 3.92, *p* = 3.01 x 10^-3^). Further, this difference appeared for all six pairs of regions and was significant for four of them (**Figure 7c**). The only non-significant increases between individual regions were right IFG to right IPS (difference in *r* values = .036, *t*(44) = 1.98, *p* = .054) and right IFG to left IFG (difference in *r* values = .037, *t*(44) = 1.70, *p* = .096). Conversely, the connectivity between left cerebellum to the frontoparietal regions showed no change from Block 1 to Block 2 (average difference in *r* values = .002, *t*(44) = .117, *p* = .907). Thus, task learning specifically increased the connectivity within the four frontoparietal regions, but not between the frontoparietal regions and the cerebellum.

**Figure 7.**
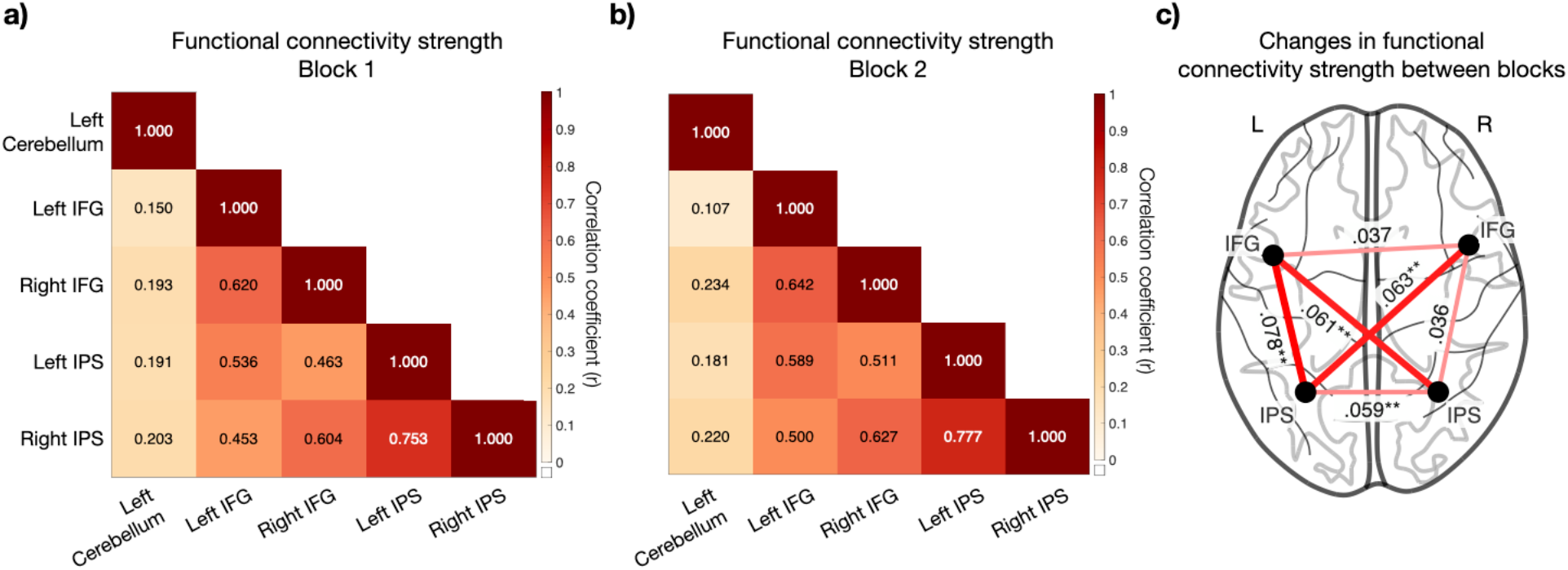
Functional connectivity for Blocks 1 and 2 between the five ROIs. (a) Average region-to-region functional connectivity for Block 1. (b) Average region-to-region functional connectivity for Block 2. (c) Change in functional connectivity strength between the two blocks. Warm colors indicate increased connectivity in Block 2 compared to Block 1. Both the thickness of the edges and the numerical values represent the change in connectivity from Block 1 to Block 2. \*\**p* < .01

## Discussion

We investigated the extent to which task learning is subserved by domain-general vs. domain-specific neural mechanisms. We found strong evidence for the domain-general account: the same five frontal, parietal, and cerebellar regions subserved the learning of six disparate tasks, while virtually no domain-specific regions were found to be involved. Further, task learning increased the functional connectivity between the frontal and parietal areas, suggesting altered communication between these areas. These results demonstrate that unlike the learning of specific stimuli, movements, or rewards, the learning of an entirely new task is domain-general and thus may underlie the human ability for acquiring new abilities.

Most previous research on how humans learn has focused on specific sensory stimuli (Antzoulatos & Miller, 2011; Rainer et al., 2004; Yang & Maunsell, 2004), motor movements (Bassett et al., 2015; Grafton et al., 2002; Houweling et al., 2008; Musall et al., 2019), and rewards (Baeuchl et al., 2020; Serences, 2008; Shuler & Bear, 2006; Summerfield & Koechlin, 2010). These studies found that each area of learning is subserved by domain-specific brain areas, making it seem like all learning – including for entirely new tasks – depends on domain-specific regions.

A growing literature has examined the neural correlates of task learning (for reviews, see Cole et al., 2013, 2017 and Meiran et al., 2017). Thus far, this literature has focused on studying different task instances drawn from the same overarching task meta-structure. In some studies, each new task involves a different type of stimulus-response mapping (Cohen-Kdoshay & Meiran, 2009; Dumontheil et al., 2011; Hartstra et al., 2011; Meiran et al., 2015a, 2015b; Muhle-Karbe et al., 2017; Ruge & Wolfensteller, 2010). In other studies, subjects first learn several elementary rules and then perform a variety of tasks where the elementary rules are combined in novel ways (Cocuzza et al., 2020; Cole et al., 2010; Cole, Reynolds, et al., 2013; Stocco et al., 2012). This line of research has identified different frontoparietal areas as critical for implementing a new task instruction. However, because in these studies each task is a specific instance of a previously learned meta-structure, the neural substrates of learning tasks with completely different meta-structures remain unclear.

To the best of our knowledge, the current study represents the first investigation of learning multiple new tasks that rely on different underlying processes. Our six tasks required the involvement of perceptual, motor, or various cognitive (attention, expectation, speed-accuracy tradeoff, and metacognition) processes. As such, subjects had to learn different task meta-structures and not just specific task instances from a learned meta-structure. By providing strong evidence for domain-specific substrates among these six tasks, our results suggest that learning novel task structures is subserved by a domain-general frontoparietal network consisting of bilateral IFG and bilateral IPS.

The frontoparietal areas identified here have partial overlap with several intrinsic brain networks that have been identified with resting fMRI data (Menon & D’Esposito, 2021), including the fronto-parietal, ventral attention, and dorsal attention networks (Chand et al., 2017; Power & Petersen, 2013; Ptak et al., 2017; Vossel et al., 2014). These networks are generally important for goal-oriented, cognitively demanding tasks, as well as for both maintaining and manipulating information (Asplund et al., 2010; Causse et al., 2022; Dixon et al., 2018; Naghavi & Nyberg, 2005). Similarly, the areas identified here have partial overlap with the nodes of a network labeled “the multiple demand network” which is associated with the execution of diverse cognitive operations (Duncan, 2010; Fedorenko et al., 2013). However, despite the partial overlap with previously identified networks, the task-learning regions we uncovered are do not map onto a single known network. Future work should determine if the task-learning network reflects a reconfiguration of known intrinsic networks or constitutes a unique task network.

The current study has several limitations. While our results constitute a critical step in establishing the domain generality of task learning, they reveal little about the functions of the specific brain areas identified here (bilateral IFG, bilateral IPS, and left cerebellum). For example, we found that the left cerebellum has weak connectivity to the remaining four frontoparietal areas, and is thus likely to have a different function than the remaining regions. It is possible that the cerebellum is involved in learning how to transform internal variables into motor commands, but it is also possible that it engages in other high-level cognitive functions that are not specifically motor-related (Buckner, 2013; Strick et al., 2009). Similarly, both the bilateral parietal and frontal areas identified here have been associated with a large array of high-level functions from attention, to working memory, to cognitive control (Corbetta & Shulman, 2002; Power & Petersen, 2013; Ptak et al., 2017; Vossel et al., 2014), and therefore it is difficult at this point to isolate the cognitive processes that these areas are likely to subserve in the context of task learning. Follow-up studies should therefore specifically focus on determining the precise function that each area is performing.

## Methods

### Subjects

Forty-eight subjects participated in the study. Data from three subjects were excluded due to data saving errors. All analyses were conducted on the remaining 45 subjects (25 females, age = 23.2±8.28 years, mean±SD). All subjects had normal or corrected-to-normal vision, were righthanded, and had no history of neurological disorders. The subjects were screened for MRI safety and provided informed consent. The study was approved by the local review board.

### Tasks

Subjects performed six different tasks that required the engagement of perception, various cognitive (attention, expectation, SAT, and metacognition), or motor processes (Figure 1). The perception task was always learned first, followed by the four cognitive tasks in a randomized order, and the motor task that always came last. Each task had three variants (see below). Each task variant consisted of two identical blocks of 30 trials (for a total of 60 trials per variant for each task). Subjects encountered each task inside the scanner for the first time and received brief on-screen instructions at the beginning of each task, as well as before the start of each new variant. Subjects were allowed unlimited time to read and process the instructions and had a 9.5-second break between the two blocks of a single task variant. Subjects gave their responses through an MRI-compatible button box with their right hand.

#### Perception task

The perception task required subjects to engage in challenging perceptual discrimination without the need to employ higher cognitive processes. Subjects indicated the orientation of a grating that could be tilted either counterclockwise (“left”) or clockwise (“right”) of vertical (Figure 1a). Each trial started with a blank screen (500 ms), followed by a white fixation point (500 ms), the grating (50 ms), and an untimed response period. The grating stimuli for the three task variants differed in their size (2°, 1.5°, and 3° in diameter), noise-level (50%, 10%, and 30%), spatial frequency of the grating (9, 3, and 20 cycles per degree), and offset from vertical (10°, 45°, and 25°). The contrast of the grating was initially set to 10% in the beginning of each task variant and was then continuously adjusted using 2-down-1-up staircase method (step size = 2% contrast) with the lowest possible contrast set to 2%. The stimuli for the four cognitive tasks were identical to the stimulus in variant 3 of the perception task, but with a fixed contrast equal to the mean contrast values of all trials in Block 2 of variant 3.

#### Attention task

The attention task required subjects to engage in the deployment of endogenous spatial attention (Carrasco, 2011; Chun et al., 2010; Petersen & Posner, 2012; Rahnev, Maniscalco, et al., 2011). Subjects saw between 2 and 4 gratings positioned 5° from the center of the screen and equally spaced from each other. On each trial, one of the gratings was post-cued and the task was to indicate the orientation of the post-cued grating (Figure 1b). A trial consisted of a blank screen (500 ms), followed by a pre-cue (an arrow) that pointed to the likely location of the post-cued stimulus (500 ms). After a fixation screen of variable duration (200-700 ms), the gratings were presented (50 ms), and finally a post-cue appeared indicating which grating subjects should respond to. Subjects had unlimited amount of time to indicate the orientation (left or right) of the post-cued grating. The three variants differed in the number of gratings presented (2, 3, or 4) as well as the predictiveness of the pre-cue (67%, 50%, and 40%), which was chosen such that the cued location was always two times more likely to be post-cued than the remaining locations.

#### Expectation task

The expectation task required subjects to engage in the integration of non-perceptual knowledge (in the form of a prior) and perceptual information (Bang & Rahnev, 2017; Rahnev, Lau, et al., 2011; Summerfield & De Lange, 2014; Turk-Browne et al., 2010). Subjects made a perceptual decision combining the information of a predictive cue with the perceptual information from the actual stimulus (Figure 1c). Each trial began with a blank screen (500 ms), followed by the predictive cue consisting of the words “LEFT” or “RIGHT” shown at the center of the screen (500 ms). After a blank screen of variable duration (200-700 ms), the grating was presented (50 ms) and was followed by an untimed response period. The three variants differed in the predictiveness of the cue, which was set to 83.3%, 95%, and 76.6%, respectively. Subjects were fully informed about the cue predictiveness and encouraged to take the information into account when making their perceptual decision.

#### Speed-accuracy tradeoff (SAT) task

The SAT task required subjects to flexibly adjust the speed of their responses (Drugowitsch et al., 2015; Giordano et al., 2009; Steinhauser & Yeung, 2012). Subjects made a perceptual decision that emphasized speed vs. accuracy to a different degree in accordance with a preceding cue (Figure 1d). The three variants differed in the number of possible instructions of the cue (variant 1: Fast/Accurate; variant 2: Fast/Neutral/Accurate; variant 3: Very fast/Fast/Neutral/Accurate). Each trial began with a blank screen (500 ms), followed by a cue that presented an instruction word at the center of the screen (500 ms). After a blank screen of variable duration (200-700 ms), the grating was presented and remained on the screen until subjects provided their response.

#### Metacognition task

The metacognition task required subjects to engage in evaluating their confidence level about the previously made perceptual decision (Fleming & Dolan, 2012; Rahnev, 2021; Yeon et al., 2020). After making a perceptual decision, subjects rated how confident they were about the decision they just made (Figure 1e). Each trial began with a blank screen (500 ms), followed by a fixation screen (500 ms). After a short presentation of a grating stimulus (50 ms), subjects sequentially indicated their perceptual decision and then provided their confidence level (both untimed). The three variants differed in the granularity of the rating scale (2, 3, and 4 points) with later variants requiring increasingly more granular confidence responses.

#### Motor task

The motor task required subjects to perform a sequence of simple button presses that required minimal involvement of higher cognitive functions (Debas et al., 2010; Doyon et al., 2018; Grafton et al., 2002). Subjects were asked to press buttons corresponding to the digits presented on the screen (Figure 1f). Each trial began with a blank screen (500 ms), followed by a stimulus presentation screen. The variants differed the number of digits presented on the screen (3, 5, and 7 digits). Numbers between one to four were used to generate a series of digits. Subjects used the index, middle, ring, and little fingers of their right hand to press buttons associated with the numbers 1-4, respectively. Subjects were instructed to perform the button presses as fast and accurately as possible.

### Image acquisition and preprocessing

The MRI data were collected on a 3T MRI system (Trio Tim, Siemens) using a 12-channel head coil. Anatomical images were acquired using T1-weighted magnetization-prepared rapid acquisition gradient-echo (MPRAGE) sequences (TR = 2,300 ms; TE = 2.98 ms; 160 slices; FoV = 256 mm; flip angle = 9°; voxel size = 1.0 x 1.0 x 1.0 mm^3^). Functional images were acquired using T2*-weighted gradient echo-planar imaging (EPI) sequences (TR = 2,000 mm; TE = 24 ms; 37 slices; FoV = 224 mm; flip angle = 60°; voxel size = 3.5 x 3.5 x 4.2 mm^3^). After subjects were positioned inside the MRI scanner, functional images were acquired continuously in a single run, during which subjects read the instructions for and performed all six tasks. The anatomical images were collected at the end of the experiment.

We used SPM12 (Wellcome Department of Imaging Neuroscience, London, UK) for data preprocessing and analysis. Functional images were first converted from DICOM to NIFTI. Preprocessing was conducted with following steps: de-spiking, slice time correction, realignment, coregistration, segmentation, normalization, and smoothing with 6-mm full-widthhalf-maximum (FWHM) Gaussian kernels.

### Data analysis

#### Behavioral analyses

We computed task accuracy and RT for each block of each task variant. For the perception and the four cognitive tasks, task accuracy was computed based on whether subjects correctly identified the grating orientation. For the motor task, a response was only considered correct if all digits were entered correctly. The RT for the motor task was computed as the time from digit presentation until the final digit was entered. We excluded outlier RT values that lie ±3 *SD* of all RTs for a given task variants. We examined differences in RT between Blocks 1 and 2 within each variant using one-sample *t*-tests, and compared these differences across task variants using paired-sample *t*-tests. Additional analyses for the four cognitive tasks, intended to confirm that each task was completed appropriately, are described in Supplementary Figures 1-4.

#### General linear model (GLM) analyses of the fMRI data

To reveal the brain regions that reflect task learning, we compared the blood-oxygenation level-dependent (BOLD) signal between Blocks 1 and 2 of each task variant. We first defined GLM regressors for Block 1 (all tasks and variants together), Block 2 (all tasks and variants together), the rest period between Blocks 1 and 2, and the instructions period that occurred for every new task and variant. The contrast Block 1 > Block 2 from this GLM was used to identify the brain areas that reflect task learning. In addition, we created two more GLMs where the periods for Block 1 and Block 2 were defined separately for each task, or separately for each task variant. The contrast Block 1 > Block 2 from these GLMs was used to examine how the brain activations differed across tasks and task variants. All GLMs included six regressors related to head movement (three translation and three rotation regressors), four tissue regressors (white matter, cerebrospinal fluid and bone, soft tissues, and air and background), and a constant term. Unless otherwise specified, analyses were performed using familywise error (FWE) corrected *p* < .05 and a cluster size of at least 150 voxels.

#### ROI analyses

To examine the domain generality of the brain regions that reflect task learning, we followed the following procedure. For a given task, we examined the contrast Block 1 > Block 2 using the data from the remaining five tasks. We then identified each of the five ROIs (bilateral IFG, bilateral IPS, and left cerebellum) based on the activations obtained using the data from these five tasks. This procedure avoided “double-dipping” where the same data are used both to define an ROI and examine its associated activations (Kriegeskorte et al., 2009). Finally, we tested whether activity in these ROIs was stronger for Block 1 than Block 2 in the original task, with the same procedure repeated for each of the six tasks.

#### Functional connectivity analyses

Our main analyses revealed that bilateral IFG, bilateral IPS, and left cerebellum reflect task learning, but could not uncover whether task learning affected the communication between these regions. To address this question, we defined these clusters as ROIs, and then extracted and normalized the time-series data from the preprocessed functional images for each ROI. We regressed out the mean gray matter, mean white matter, and mean cerebrospinal fluid signals from the time-series for each ROI (Ciric et al., 2017; Lydon-Staley et al., 2019). We also removed any individual volumes with framewise displacement greater than 0.3 mm, to minimize artifacts related to in-scanner head motion (Jenkinson et al., 2002; Power et al., 2012). Lastly, we filtered the residual signal with a second-order Butterworth filter between 0.01-0.1 Hz. Finally, we computed the strength of the functional connectivity between each pair of the five ROIs separately for Blocks 1 and 2. To statistically compare the correlation values between Blocks 1 and 2, we z-transformed the *r*-values and conducted paired-sample *t*-tests on the *z*-values. All average *r*-values reported in the main text and Figure 7 were obtained by averaging the *z*-values and then transforming the average *z*-value back into an *r*-value.

### Data and code

Data and analysis code are available at https://osf.io/knzj6. In addition, all group-level fMRI T-maps are available at https://neurovault.org/collections/FYJMWHCU.

## Acknowledgment

This work was supported by the National Institute of Mental Health of the National Institutes of Health under Award Number R01MH119189 to D.R.

## Competing Interest Statement

The authors declare no competing interests

## Notes

### Competing Interest Statement

The authors have declared no competing interest.

