## Supplementary materials for "Task learning is subserved by a domain-general brain network"

**A domain-general brain network reflects the learning of new tasks**

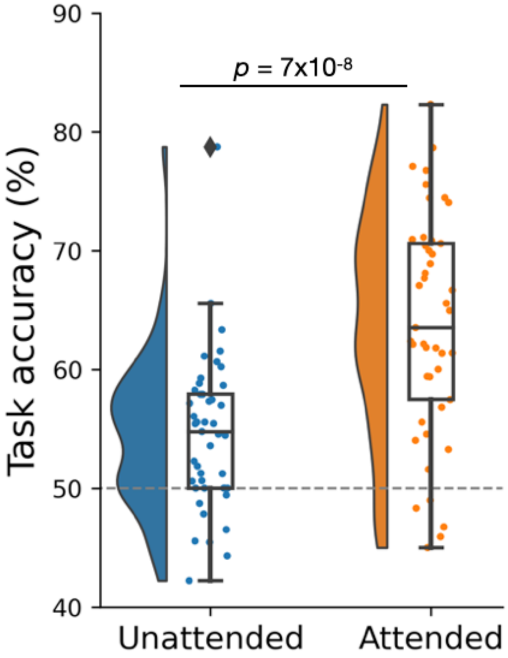


**Supplementary Figure 1. Manipulation check for the attention task**. Subjects who follow the attention instructions would be expected to have higher performance for the attended targets (the targets that were pre-cued) than for the unattended targets (the targets that were not pre-cued) (Rahnev et al., 2011). Indeed, subjects showed higher task performance when the target was attended (average accuracy = 63.8%, SD = 9.4%) than when it was unattended (average accuracy = 54.5%, SD = 6.5%; $t(44) = 6.46$, $p = 7 \times 10^{-8}$). These results demonstrate that subjects followed the attention cue and thus performed the task as intended. Dots represent individual subjects, and the boxes indicate interquartile range (IQR) while the inner line of the boxes shows the median value. The data points outside the whiskers of the boxes (±1.5 x IQR) are outliers. The figure was generated using Raincloud Plots (Allen et al., 2021).

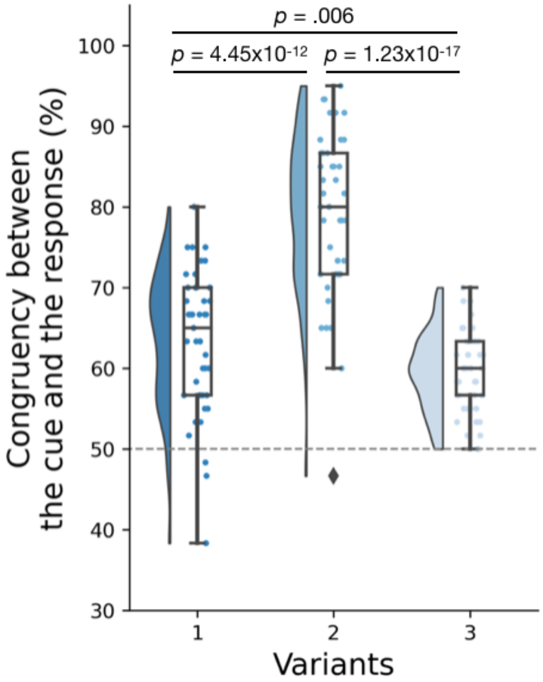


**Supplementary Figure 2. Manipulation check for the expectation task**. Subjects who follow the expectation cue would be expected to have higher-than-chance congruency between the cue and their response. In addition, the congruency should be higher for cues of higher predictiveness (Rahnev & Denison, 2018). The three task variants presented cues with validity of 83.3%, 95%, and 76.6 %, respectively, and therefore the congruency between the cue and response would be expected to be highest for the second variant and lowest for the third variant. This is exactly what we found: the congruency between the cue and the response was highest for the second variant (*M* = 79%, *SD* = 1%), followed by the first (*M* = 63.4%, *SD* = 8.6%) and the third variant (*M* = 59.5%, *SD* = 4.9%; $F(2, 132) = 72.63$, $p = 5.35 \times 10^{-22}$). These results demonstrate that subjects followed the expectation cue according to its validity and thus performed the task as intended. The box plots and the dots follow the same scheme as Supplementary Figure 1.

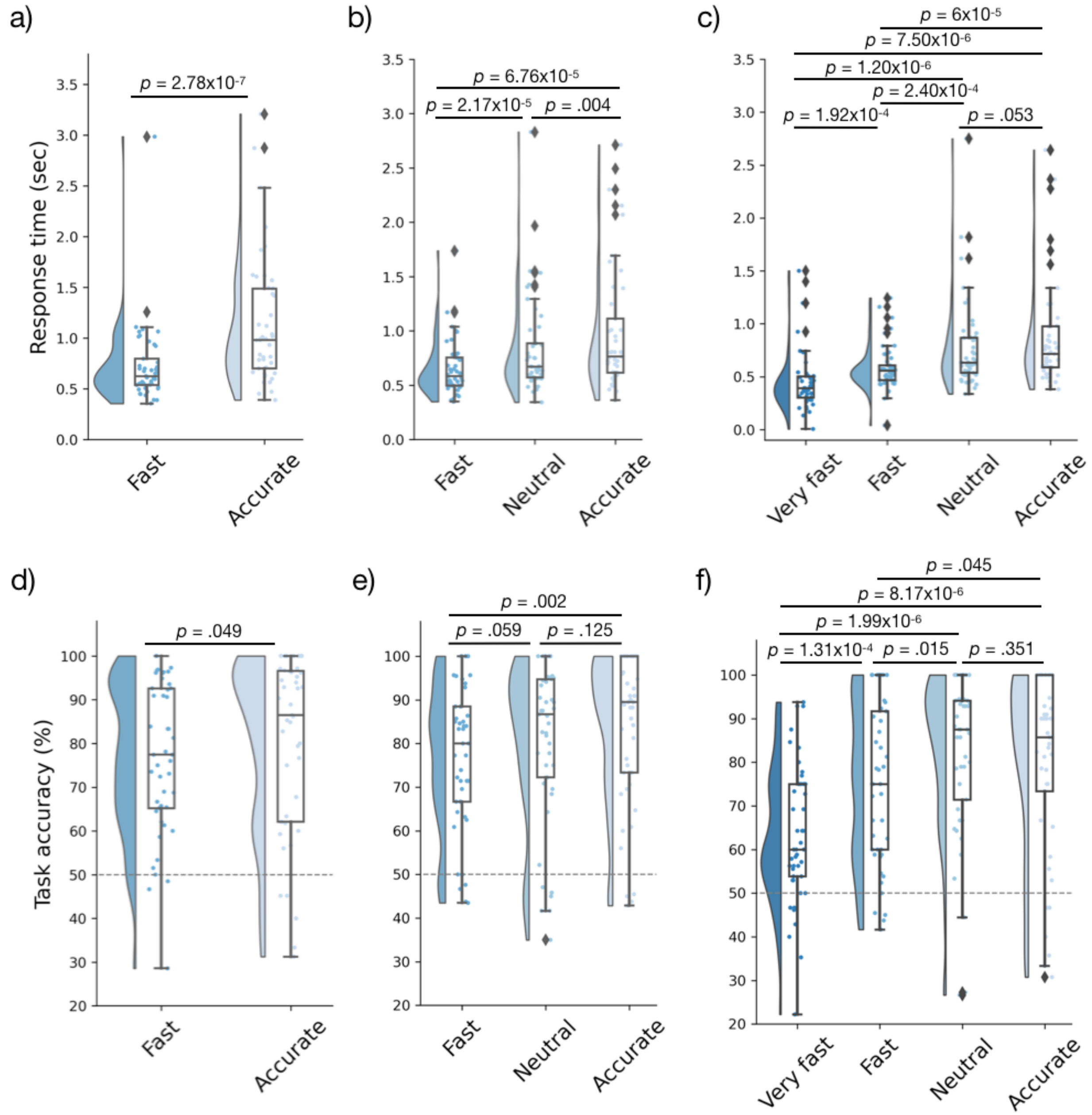


**Supplementary Figure 3. Manipulation check for the SAT task**. Subjects who follow the SAT cue would be expected to have faster and less accurate responses for higher speed stress cues. In general, these results tend to be more robust for RT compared to accuracy (Rafiei & Rahnev, 2021; Rahnev et al., 2016), which is also what we found here. (a) RT results for the first variant: Subjects responded faster when the cue 'Fast' was presented ($M_{RT}$ = .734 sec, *SD* = .409) compared to when the cue 'Accurate' ($M_{RT}$ = 1.142 sec, *SD* = .629; $t(44) = 6.057$, $p = 2.78 \times 10^{-7}$). (b) RT results for the second variant: Subjects responded faster for cues that asked them to emphasize speed more ('Fast' cue: $M_{RT}$ = .66 sec, *SD* = .264; 'Neutral' cue: $M_{RT}$ = .844 sec, *SD* = .461; 'Accurate' cue: $M_{RT}$ = .994 sec, *SD* = .581), with the difference between conditions being significant ($F(2,132) = 6.11$, $p = .003$). (c) RT results for the third variant: Subjects responded faster for cues that asked them to emphasize speed more ('Very fast' cue: $M_{RT}$ = .455 sec, *SD*

= .297; 'Fast' cue: $M_{RT}$ = .583 sec, $SD$ = .217; 'Neutral' cue: $M_{RT}$ = .781 sec, $SD$ = .432; 'Accurate' cue: $M_{RT}$ = .892 sec, $SD$ = .525), with the difference between conditions being significant ($F$(3,176) = 11.562, $p$ = 5.917 x $10^{-7}$). (d) Accuracy results for the first variant: Subjects responded less accurately when the cue 'Fast' was presented ($M_{accuracy}$ = 76.3%, $SD$ = 17.3%) compared to when the cue 'Accurate' was presented ($M_{accuracy}$ = 80%, $SD$ = 20.3%; $t$(44) = 2.02, $p$ = .049). (e) Accuracy results for the second variant: Subjects responded less accurately for cues that asked them to emphasize speed more ('Fast' cue: $M_{accuracy}$ = 76.4%, $SD$ = 16.7%; 'Neutral' cue: $M_{accuracy}$ = 80.4%, $SD$ = 18.4%; 'Accurate' cue: $M_{accuracy}$ = 83.1%, $SD$ = 17.5%). However, only the comparison between 'Fast' and 'Accurate' cues was statistically significant ($t$(44) = 3.30, $p$ = .002). (f) Accuracy results for the third variant: Subjects generally responded less accurately for cues that asked them to emphasize speed more ('Very fast' cue: $M_{accuracy}$ = 63.1%, $SD$ = 15.8%; 'Fast' cue: $M_{accuracy}$ = 75%, $SD$ = 18.4%; 'Neutral' cue: $M_{accuracy}$ = 81.9%, $SD$ = 18.6%; 'Accurate' cue: $M_{accuracy}$ = 80.1%, $SD$ = 20.7%), with the difference between the different cues being significant $F$(3,176) = 9.51, $p$ = 7.48 x $10^{-6}$). Overall, these results demonstrate that subjects followed the SAT cue and thus performed the task as intended. The box plots and the dots follow the same scheme as Supplementary Figure 1.

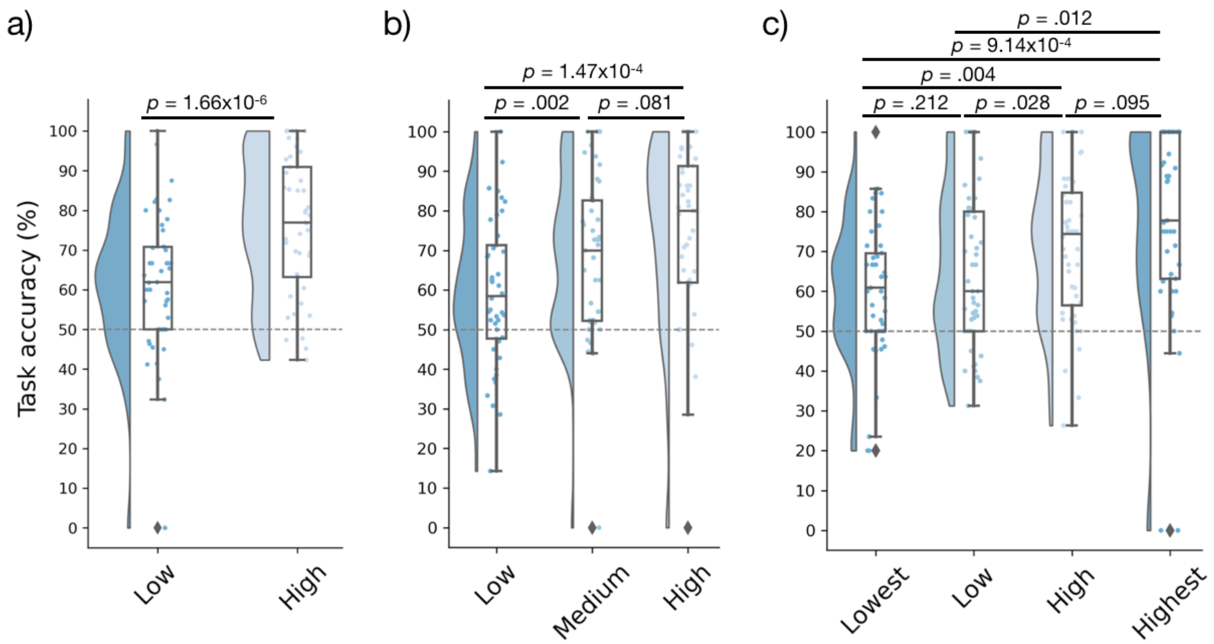


**Supplementary Figure 4. Manipulation check for the metacognition task**. Subjects who provide confidence ratings appropriately would be expected to have higher accuracy when they are more confident and lower accuracy when they are less confident (Rahnev, 2021). Indeed, we found gradually increasing task performance for higher confidence ratings for the first variant (accuracy for low confidence responses: $M$ = 61.8%, $SD$ = 17.9%; accuracy for high confidence responses: $M$ = 76%, $SD$ = 17.5%; $t(44) = 5.53$, $p = 1.66 \times 10^{-6}$), for the second variant (accuracy for low confidence responses: $M$ = 59.4%, $SD$ = 18.8%; accuracy for medium confidence responses: $M$ = 68.2%, $SD$ = 20.4%; accuracy for high confidence responses: $M$ = 73.3%, $SD$ = 24.5%; $F(2, 131) = 4.80$, $p = .01$), and for the third variant (accuracy for lowest confidence responses: $M$ = 60.2%, $SD$ = 17.2%; accuracy for low confidence responses: $M$ = 64.7%, $SD$ = 19.1%; accuracy for high confidence responses: $M$ = 71.2%; $SD$ = 18.8%; accuracy for highest confidence responses: $M$ = 76.1%, $SD$ = 26.6%; $F(3, 175) = 5.09$, $p = .002$). These results demonstrate that subjects provided appropriate confidence ratings and thus performed the task as intended. The box plots and the dots follow the same scheme as Supplementary Figure 1.

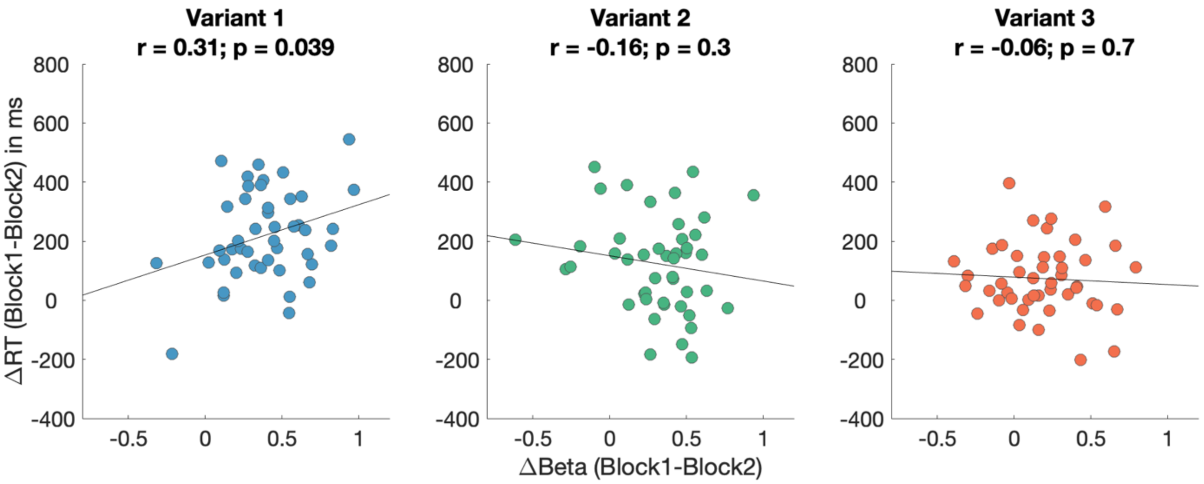


**Supplementary Figure 5. Across-subject brain-behavior correlations for each task variant**. We correlated the RT difference between Blocks 1 and 2 (normalized within tasks and versions, and averaged across the six tasks) and the beta value difference between Blocks 1 and 2 (averaged across all five ROIs: bilateral MFG, bilateral precuneus, and left cerebellum). We found a significant correlation for the first variant ($r$ = .308, $p$ = .039), but no such correlation for the second ($r$ = -.157, $p$ = .303) and third task variants ($r$ = -.059, $p$ = .702). These results suggest that subjects who showed more behavioral learning also exhibited larger brain responses in the five frontal, parietal, and cerebellar regions, but that this effect only holds for the first task variant where the majority of learning takes place. Nevertheless, such across-subject brain-behavior correlations should be interpreted with caution when performed at the sample sizes used in our current study (Marek et al., 2022).
